# The extent of myeloid skewing in blood is a biomarker of biological aging in mice and humans

**DOI:** 10.64898/2026.01.20.700344

**Authors:** Julian Niemann, Selina Stahl, Vadim Sakk, Juan-Felipe Perez-Correa, Wolfgang Wagner, Andrea Jaensch, Dietrich Rothenbacher, Yi Zheng, Medhanie A Mulaw, Hartmut Geiger, Mona Vogel

**Author notes:** Correspondence: Mona Vogel, Ulm University, 89081 Ulm, Germany. **Data availability**: Primary data will be available upon reasonable request.

## Abstract

Myeloid skewing is a central and therefore often cited hallmark of hematopoietic aging. Myeloid skewing refers to an elevated myeloid-to-lymphoid cell ratio in aged compared to young mice. Interestingly, whether the extent of myeloid skewing might be in itself a quantitative biological marker of aging has not been addressed yet, nor whether this parameter has also relevance for the extent of aging in humans. Aged mice with high level of myeloid skewing (>50% myeloid cells in blood) showed accelerated hematopoietic aging compared to mice with a low level of myeloid skewing (<30% of myeloid cells in blood), as well as an increased level of inflammatory cytokines and elevated levels of diseases. Hematopoietic stem cells (HSCs) from mice with high myeloid skewing showed an impaired repopulation capacity. Epigenetic clock analyses demonstrated that mice with a high level of myeloid skewing present with a biological age that is older than their chronological age. In humans, a high degree of myeloid skewing was associated with elevated levels of inflammatory markers, reduced mobility, a greater burden of comorbidities, and an increased mortality hazard ratio. The data support that, besides overall myeloid skewing being a central hallmark of aging in mice, the extent of the frequency of myeloid cells in blood might serve as a biological marker of aging and disease in both mice and humans.

**Key Points:** The extent of myeloid skewing in aged mice correlates to an increased hematological and epigenetic age and increased disease burden.

The extent of myeloid skewing in older adults is associated with an increased hazard ratio of mortality and correlates with higher frailty and inflammatory markers.

## INTRODUCTION

The rate of aging varies significantly among individuals, even among those with a similar chronological age. This inter-individual variability resulted in the concept of biological age, a parameter to more accurately reflect the functional state the biological system of an individual rather than using its chronological age ^1–4^. Aging is, among others, accompanied by a decline in immune competence, referred to as age-associated immune remodeling^5,6^. A key change in the composition of the immune system upon aging is an increased proportion of myeloid cells (e.g., monocytes, granulocytes, dendritic cells and megakaryocytes) relative to lymphoid cells (e.g., B and T lymphocytes), an observation known as myeloid skewing^7–10^. The increased relative proportion of myeloid cells has been linked to a chronic low-grade inflammation in older organism ^11–13^. Such a low-grade inflammatory milieu might be a central driver of systemic aging^14,15^.

In mice, myeloid skewing is a well-established hallmark of hematopoietic aging. Myeloid skewing arises from a change in the differentiation potential of hematopoietic stem cells (HSCs)^16–19^. Young HSCs show a balanced differentiation into myeloid and lymphoid cells, whereas aged HSCs increasingly favor the production of myeloid over lymphoid cells^8–10,16^. Interestingly, information on the dynamics, the stability and extent of myeloid skewing in individual animals over time is not available^20^. In addition, in humans myeloid skewing has been reported to be much less pronounced, while its potential as aging-marker has not yet been established^21^.

We demonstrate here that the extent of myeloid skewing is a robust quantitative biomarker for aging of the hematopoietic compartment for both murine and human aging, with a strong correlation to inflammation, disease and general aging-associated health parameters as well as mortality in humans. This finding supports the use of the extent of myeloid skewing as a readout for pre-clinical aging studies in animals.

## METHODS

### Mice

In all experiments, female C57BL/6J mice were used. Male and female B6.SJL-Ptprca Pepcb/BoyJ mice served as recipients in the transplantation experiments. All animals were housed and bred under specific pathogen-free conditions at the animal housing facility at Ulm University or the Association for Assessment and Accreditation of Laboratory Animal Care-accredited animal facility of Cincinnati Children’s Hospital Medical Center (CCHMC). All work done on animals was in accordance with the protocols from the institutional animal care and use committee at Cincinnati Children’s Hospital Medical Center and the German Law for Welfare of Laboratory Animals in Tübingen. Aging cohorts (>80 weeks) were bled and classified into aged^low^ and aged^high^ groups. For tissues analyses, mice were selected from these classified groups according to similar age and, whenever possible, housed in the same cage to minimize external influences.

### HSC Isolation and Transplantation

On the day of transplantation, myeloid percentages were re-verified in old donor mice (female, C57/BL6J, >80 weeks; CD45.2^+^) and mice were screened for cross-tissue abnormalities. Mice displaying a changed in the percentage of myeloid skewing compared to the pre-bleeding or pathological changes in more than one hematopoietic or hematopoietic-associated tissue were excluded from providing donor cells for transplantation experiments. Bone marrow (BM) was isolated from young (female, C57/BL6J, 12-16 weeks; CD45.2^+^) and old mice by flushing of bones. Mononucleated cells were isolated with a low density Ficoll (Cytiva) gradient. Lineage depletion was used to enrich the primitive cell population (Miltenyi). The resulting cells were stained with antibodies and HSCs (Lin^-^, Sca-1^+^, c-Kit^+^, CD34^-^, Flt3^-^) were isolated with FACS using an Aria III (BD Biosciences). For transplantation experiments, 500 HSCs were isolated and intravenously injected into irradiated (7+4 Gy) recipient mice (B6.SJL-Ptprc^a^ Pepc^b^/BoyJ, CD45.1^+^, 8-14 weeks) together with 10^5^ total BM competitor cells (B6.SJL-Ptprc^a^ Pepc^b^/BoyJ, CD45.1^+^). Transplanted mice were bled every 4 weeks and PB was analyzed for engraftment and lineage contribution. 16 weeks after transplantation mice were sacrificed and peripheral blood (PB), spleen and BM analyzed.

### Organ analysis

PB was analyzed using a ScilVet abc Plus (ScilVet) for complete blood counts, erythrocyte, and platelet parameters, and a LSRFortessa flow cytometer (BD Biosciences) was used to characterize different cell subtypes by flow cytometric analyses. Spleens were passed through a 70 µm cell strainer to obtain single-cell suspensions, and BM was isolated by flushing the femurs, tibias, and hip bones of the mice. PB, spleen and BM were subjected to red blood cell lysis (Biolegend) prior to antibody staining. Hematological cell populations in PB, BM and spleen were stained with a combination of CD45.2, CD11b, Gr-1 CD3/CD4/CD8, CD19, CD44 and CD62L. Splenocytes were additionally stained with CD45.2, CD19, CD21/32 and CD23. Finally, we analyzed lineage depleted BM for primitive cells using different combinations of CD45.2, Lin, Sca-1, c-Kit, CD34, CD150, CD48, CD16/32 and IL7Ra.

### Cytokine measurement

To obtain blood serum, mice were sacrificed and blood was collected by cardiac puncture. Samples were kept for 20–30 minutes at room temperature in microvette tubes containing serum gel with a clotting activator (Sarstedt, Ref. 20.1344). Blood was then centrifuged, and the serum was collected, aliquoted, and stored at −80 °C until further use. BM was isolated from two femurs and two tibias per mouse, centrifuged into 50 µL of PBS, resuspended, centrifuged again, and the resulting supernatant was collected and stored at −80 °C until further processing.

The Mouse Anti-Virus Response Panel Mix and Match Kit was used (Biolegend, Lot Standard: B332105) and was performed according to the manufacturer’s instructions for V-Bottom plates with the following modifications: serum and BM supernatant samples were applied undiluted, and half of the recommended reagent volumes were used for all steps. Standards were prepared as described in the manual. All samples were run in duplicates. The following cytokines were measured: MCP-1, KC, IL-10, CCL5, IL-6, IFN-γ, and IL-1β. Data acquisition was performed using a Cytek Aurora flow cytometer (Cytek Biosciences), and setup and analysis were carried out according to the Biolegend protocol.

### RNA Sequencing of hematopoietic stem cells

Murine HSCs (LSK, CD34^low^, Flt3^-^) were sorted and processed according to the SMART Seq. v4 Ultra Low Input RNA Kit (Takara Bio). Libraries were prepared with the Nextera XT library Kit (Illumina). Raw data were processed, aligned and mapped as previously described^5,22^.

### Epigenetic analysis of peripheral blood

Blood of young, aged^low^ and aged^high^ mice was collected by cardiac puncture and stored in EDTA microvette tubes (Sarstedt) at -80°C. DNA methylation levels were assessed at four age-associated CpG sites as previously described^23^. Briefly, genomic DNA was extracted from the whole blood samples, subjected to bisulfite conversion, and analyzed by pyrosequencing for methylation at four genes: *Hsf4*, *Prima1*, *Aspa*, and *Wnt3a*. These methylation values were then incorporated into a multivariable model^23,24^ for predicting epigenetic age in C57BL/6 mice, which has been shown to correlate strongly with chronological age. To assess the impact of myeloid/lymphoid cell proportions on the age predictions, methylation data from nine distinct FACS-sorted murine hematopoietic cell types, each represented by biological triplicates, was retrieved from the Gene Expression Omnibus (GSE201923). The aging model described previously was employed to calculate the epigenetic age of these various cell types. The analysis revealed no significant difference (p = 0.16) between the predicted ages of myeloid and lymphoid cells, which facilitated the implementation on the young, aged^low^ and aged^high^ mice samples without requiring any further modifications of the model.

### Data analysis and statistics (mouse)

GraphPad Prism (versions 9 and 10; GraphPad Software) was used for data visualization and statistical analysis. For each experiment and group, outliers were identified and removed using the ROUT method (Q = 1%). Normality was assessed using the Shapiro–Wilk test. Unless otherwise specified, statistical analyses were performed using one-way ANOVA with Tukey’s post-hoc test for normally distributed data, or Kruskal–Wallis test with Dunn’s multiple-comparison correction for nonparametric data among young, aged^low^, and aged^high^ groups. Nonparametric tests were applied when any group failed the normality test. Comparisons between young and old cohorts were performed using either the Mann–Whitney test or Welch’s t-test, depending on data distribution. Biological replicates are indicated in the respective figure legends. All experiments were independently repeated at least three times. Data are presented as medians with 95% confidence intervals (CI) unless indicated. Illustrations were created using BioRender.

### Population-based cohort: ActiFE study data analyses

Human data were retrieved from the population-based ActiFE Study, a cohort study in subjects aged 65+ years at baseline, randomly selected from Ulm and adjacent regions. Details can be found in the published study protocol and other original publications^25,26^. A standardized assessment at baseline was completed by trained research assistants. Exclusion criteria were being in residential care, having serious German language difficulties, or not able to give informed consent. The baseline study took place between March 2009 and April 2010 and N=1506 older adults participated. Blood was collected during the interview visit and baseline laboratory tests were immediately done in the routine laboratory of the Agaplesion Bethesda clinic including a complete blood cell count. All participants gave written consent. The ethical review board of Ulm University had approved the study (No. 318/08 and 50/12). All participants with data for myeloid skewing were included in the analysis (n=1425, 95% of overall population). Data were stratified into quartiles based on their extent of myeloid skewing. Stratification and descriptive characteristics of patients and measurements can be found in **Table 1**. Mortality status and date of death were obtained from the local registration offices for all participants. Cox proportional hazards models were used to estimate the association of myeloid skewing (in quartiles) with four, six, and eight year mortality after adjustment for age and sex, and also additional adjustment for well-established covariates (i.e. age, sex, school education (<=9/>9 years), body mass index (BMI) (categorised in four categories: underweight, normal, overweight, obese), current smoker, alcohol consumption (daily/less than daily), doctor diagnosed history of hypertension, myocardial infarction, cancer, diabetes, respectively, and number of medications (<5/>=5)).

**Table 1:** Baseline characteristics of study population, overall and according to the % of myeloid skewing quartiles. SD standard deviation. N corresponds to sample size.

| Characteristic | Total<br>(n=1425) | older<br>adults<sup>low</sup><br>(Q1)<br>(n=350) | Q2<br>(n=356) | Q3<br>(n=364) | older<br>adults<sup>high</sup><br>(Q4)<br>(n=355) |
| --- | --- | --- | --- | --- | --- |
| Chronological age (years), mean (SD) | 75.58<br>(6.59) | 73.62<br>(6.04) | 74.88<br>(6.38) | 75.96<br>(6.55) | 77.82<br>(6.68) |
| Women, n(%) | 614 (43.1) | 198 (56.6) | 176 (49.4) | 146 (40.1) | 94 (26.5) |
| Duration of school education ≤9 y, n(%) | 809 (57.6) | 195 (56.7) | 197 (56.1) | 198 (55.3) | 219 (62.2) |
| Current smoker, n(%) | 105 (7.4) | 29 (8.3) | 29 (8.2) | 24 (6.6) | 23 (6.5) |
| Daily alcohol consumption, n(%) | 434 (31.0) | 98 (28.4) | 92 (26.1) | 127 (35.7) | 117 (33.5) |
| BMI (kg/m<sup>2</sup>), mean (SD) | 27.60<br>(4.16) | 27.52<br>(4.12) | 27.68<br>(4.34) | 27.62<br>(4.06) | 27.58<br>(4.12) |
| ≥30 kg/m<sup>2</sup>, n(%) | 347 (24.4) | 85 (24.3) | 96 (27.0) | 83 (22.8) | 83 (23.4) |
| Mini-Mental State Examination <25 points, n(%) | 72 (5.5) | 17 (5.3) | 13 (4.0) | 15 (4.5) | 27 (8.0) |
| Self-reported doctor-diagnosed comorbidity, n(%) |  |  |  |  |  |
| Hypertension | 762 (53.7) | 173 (49.6) | 187 (52.5) | 195 (53.9) | 207 (58.6) |
| Heart failure | 207 (14.6) | 47 (13.5) | 41 (11.5) | 50 (13.9) | 69 (19.5) |
| Myocardial infarction | 126 (8.9) | 22 (6.3) | 31 (8.7) | 30 (8.3) | 43 (12.2) |
| Cancer | 261 (18.4) | 42 (12.0) | 64 (18.0) | 73 (20.2) | 82 (23.4) |
| Diabetes | 189 (13.3) | 51 (14.6) | 41 (11.5) | 41 (11.3) | 56 (15.8) |
| Osteoporosis | 152 (10.7) | 34 (9.7) | 41 (11.5) | 38 (10.5) | 39 (11.0) |
| Self-rated health “fair or bad”, n(%) | 237 (16.7) | 46 (13.3) | 54 (15.2) | 63 (17.4) | 74 (21.0) |
| Gait speed (m/s), mean (SD) | 1.04 (0.33) | 1.08 (0.31) | 1.05 (0.32) | 1.03 (0.31) | 0.99 (0.38) |
| Daily mean duration (min) of walking, mean (SD) | 104.23<br>(40.33) | 108.30<br>(38.06) | 105.67<br>(39.82) | 104.44<br>(39.45) | 98.43<br>(43.37) |
| Frailty Index, mean (SD) | 0.14 (0.09) | 0.12 (0.09) | 0.13 (0.08) | 0.14 (0.10) | 0.16 (0.10) |
| Five-chair-rise, mean (SD) | 11.27<br>(3.55) | 10.66<br>(3.19) | 11.07<br>(3.26) | 11.50<br>(4.02) | 11.84<br>(3.58) |
| CRP, mean (SD) | 3.53 (7.71) | 2.62 (6.09) | 2.60 (3.38) | 3.45 (4.72) | 5.46<br>(12.76) |
| IL-6, mean (SD) | 2.90 (4.04) | 2.25 (2.63) | 2.41 (2.95) | 2.95 (4.06) | 3.97 (5.61) |
| Lymphoid cells, mean (SD) | 1.91 (1.70) | 2.68 (3.21) | 1.97 (0.45) | 1.71 (0.38) | 1.30 (0.38) |
| Myeloid cells, mean (SD) | 4.69 (1.43) | 3.71 (0.94) | 4.31 (1.02) | 4.93 (1.08) | 5.80 (1.65) |

The comparison of myeloid skewing categories (in quartiles) with various continuous variables was analyzed for the available dataset by means of a linear regression model after adjustment for age and gender in an exploratory manner. Statistical analyses were performed with SAS 9.4 (SAS Institute, Cary, North Carolina, USA).

## RESULTS

### The extent of myeloid skewing upon aging is heterogeneous

We first determined the percentage of myeloid cells among blood cells in peripheral blood (PB) throughout the life span of individual mice. The percentage of myeloid cells among blood cells did on average increase with age among all individual mice analyzed (black line **Fig.1A**). There was some heterogeneity with respect to the trajectory of the frequency of myeloid cells among blood cells among individual mice upon aging (see area between dotted black lines, **Fig.1A**). Some mice showed only an overall minor increase in the frequency of myeloid cells during aging with some minor fluctuations. These might be due to levels of acute stress as well as acute infections. Both are known to upregulate the frequency of myeloid cells^27^, while the general increase in myeloid cells over time might be due to aging. Other aged mice showed a sudden, steep increase in PB myeloid cells, with the onset varying by individual and not predictable from earlier myeloid patterns. We also evaluated the consistency of distinct experimental approaches to determine the value “percentage of myeloid cells” in murine blood. We found a robust correlation between the percentages of granulocytes obtained using the blood counter (ScilVet) and those of myeloid cells (Mac-1 and/or Gr-1 positive cells) acquired through flow cytometry (r=0.801) (**Fig.S1A**). Consequently, quantifying the degree of myeloid skewing can be reliably performed via standard veterinary blood counters. Myeloid skewing is characterized by both a reduction in the total number of lymphoid cells in PB and a slight increase in the total number of myeloid cells (**Fig.S1B**). Importantly, myeloid skewing manifested consistently across different bleeding methods (**Fig.S1C**) and remained unaffected by the circadian rhythm (**Fig.S1D**). Taken together, there is robustness in the level of myeloid cells in blood and among the approaches to determine the extent of myeloid skewing in individual mice. Consequently, the level of heterogeneity in myeloid skewing among individual mice can be used to test for correlations to other parameters of aging.

**Figure 1.**
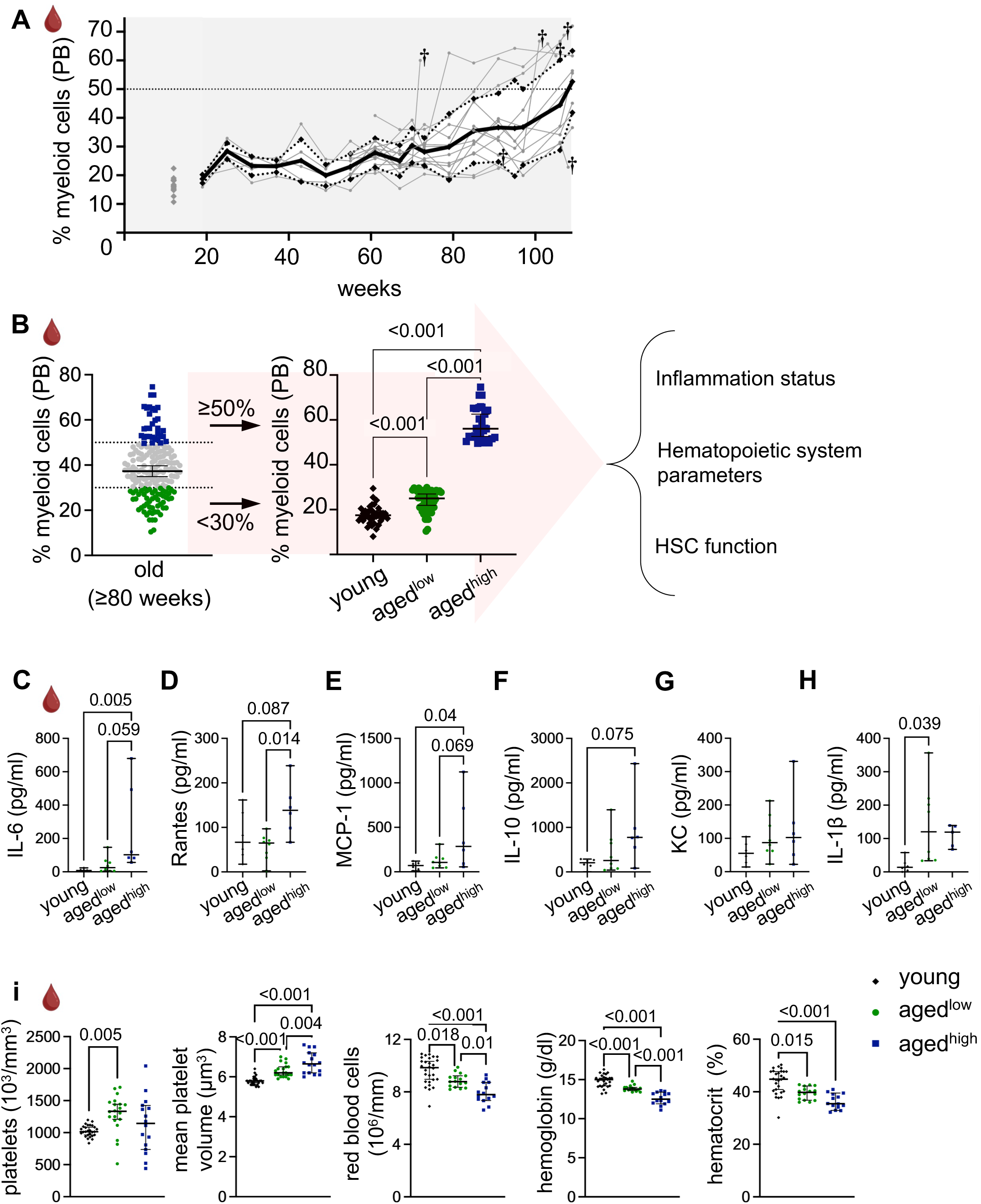
The extent of myeloid skewing upon aging is heterogeneous **A)** Longitudinal analysis of myeloid cell percentage and its variation over time in individual mice. Black line: average of extend of myeloid skewing. Black dotted line/white area: ± standard deviation. Grey lines: individual mice. Cross: Death of the mouse. n = 15 mice (start), 9 mice (end), bled every 6-8 weeks. **B)** Overview of experimental set-up: aged mice (>80 weeks) were pre-bled (n = 187) and aged^high^ (n = 32) and aged^low^ (n = 54) were used for further experiments. Young mice (10-16 weeks, n = 39) were used as controls. **C-H)** Cytokines were measured in blood serum (pg/ml). (young n = 6; aged^high^ n = 6; aged^low^ n = 8) **C)** Interleukin 6 (IL-6) **D)** C-C motif chemokine ligand 5 (Rantes) **E)** Monocyte chemoattractant protein-1 (MCP-1) **F)** Interleukin 10 (IL-10) **G)** Keratinocyte-derived chemokine (KC) **H)** Interleukin 1β (Il-1β) **i)** Platelets and Erythrocyte parameters analyzed with the blood cell counter ScilVet. (young n = 27-28; aged^high^ n = 14-16; aged^low^ n = 15-20). Data are presented as medians with 95% CI. Outliers were excluded using the ROUT method (Q = 1%). Statistical analysis was performed using one-way ANOVA with Tukey’s post-hoc test for normally distributed data or Kruskal–Wallis test with Dunn’s multiple-comparison correction for nonparametric data.

### The extent of myeloid skewing correlates with the extent of age-associated changes in the hematopoietic system

When examining aged mice (>80 weeks), we observed heterogeneity in the extend of myeloid skewing in PB. Based on this observation, we stratified the animals into aged^low^ and aged^high^ groups: the aged^low^ group exhibited low myeloid frequencies (≤30%), whereas the aged^high^ group displayed pronounced myeloid skewing (≥50%). In the analyzed cohort, 29% of mice fell into the aged^low^ group and 17% into the aged^high^ (**Fig.1B**). BM and spleen also showed increased myeloid skewing, however the overall magnitude of skewing was considerably lower than in PB (**Fig.1B/S2A**). To test whether low or high myeloid skewing correlates with other aging-associated hematopoietic parameters in mice, the cohorts were analyzed for inflammation, frequency of other hematopoietic cell populations and HSCs function (**Fig.1B**).

First, mostly aged^high^ mice—rarely aged^low^ mice—exhibited elevated levels of pro-inflammatory cytokines in PB such as interleukin-6 (IL-6), CCL5 (Rantes) and monocyte chemoattractant protein-1 (MCP-1) in blood serum when compared to young mice (**Fig.1C–E**). In addition, IL-10 (**Fig.1F**) and keratinocyte-derived chemokine (KC) (**Fig.1G**) showed a trend toward higher cytokine levels in aged^high^ compared to young mice. Together, this is indicative of a distinct type of pro-inflammatory milieu in aged^high^ mice compared to aged^low^ mice. Interestingly, not all cytokines showed this pattern, for example IL-1β was elevated between young and aged mice but not between aged^low^ and aged^high^ groups (**Fig.1H**), suggesting that only a selected set of pro-inflammatory cytokines is elevated in aged^high^ mice, thus the extend of myeloid skewing may follow a specific pattern or could be driven by specific immune cell populations whose functions may be altered upon aging. A substantial heterogeneity in cytokine levels was present within both aged^low^ and aged^high^ groups and correlation analyses between the degree of myeloid skewing and cytokine levels in blood serum of aged animals were weak (**Fig.S2B/C**). Combined with the finding that aged^low^ mice show increased myeloid frequencies without consistent cytokine elevation, this suggests that aging-associated myeloid skewing can occur independently of inflammation. However, inflammation may further exacerbate the extent of myeloid skewing, as suggested by the elevated levels of inflammation-associated cytokines in aged^high^ mice.

Second, aged^high^ animals exhibited marked alterations in platelet and red blood cell parameters (**Fig.1i**). Platelet counts were significantly increased in the aged^low^, but not the aged^high^ group in comparison to young controls. In contrast, mean platelet volume, red blood cell count, hemoglobin concentration, and hematocrit levels showed a gradual change from young to aged^low^ to aged^high^ mice, with aged^low^ animals more closely resembling young mice, indicating elevated aging-associated parameters within the erythrocyte compartment and, to a lesser extent, in platelets in aged^high^ animals.

Third, CD3⁺, CD4⁺, and CD8⁺ T-cell frequencies in PB were reduced in aged mice, even stronger in aged^high^ mice (**Fig.2A**). In the spleen, only CD8⁺ T cells declined with age (**Fig.S3A**), and no significant changes were seen in BM (**Fig.S3B**). As a reduced frequency of naïve CD8⁺ T-cells and an increase of effector memory cells upon aging are a hallmark of immune-aging, we examined CD8⁺ T-cell subsets (**Fig.2B**). As expected, we observed a substantial decline in the frequency of naïve CD8⁺ T-cells in BM, PB and spleen of aged animals and an increase in the frequency of effector memory cells (**Fig.2C/S3C/D**). In aged^high^ animals, naïve CD8⁺ T-cells tended to be further reduced and effector memory T-cells further increased relative to aged^low^ animals in both PB and spleen (**Fig.2C/S3C**). Also, for naïve CD4^+^ T-cells there was a further increase in the aging associated decrease of their frequency in aged^high^ animals in PB and spleen (**Fig.2D**).

**Figure 2.**
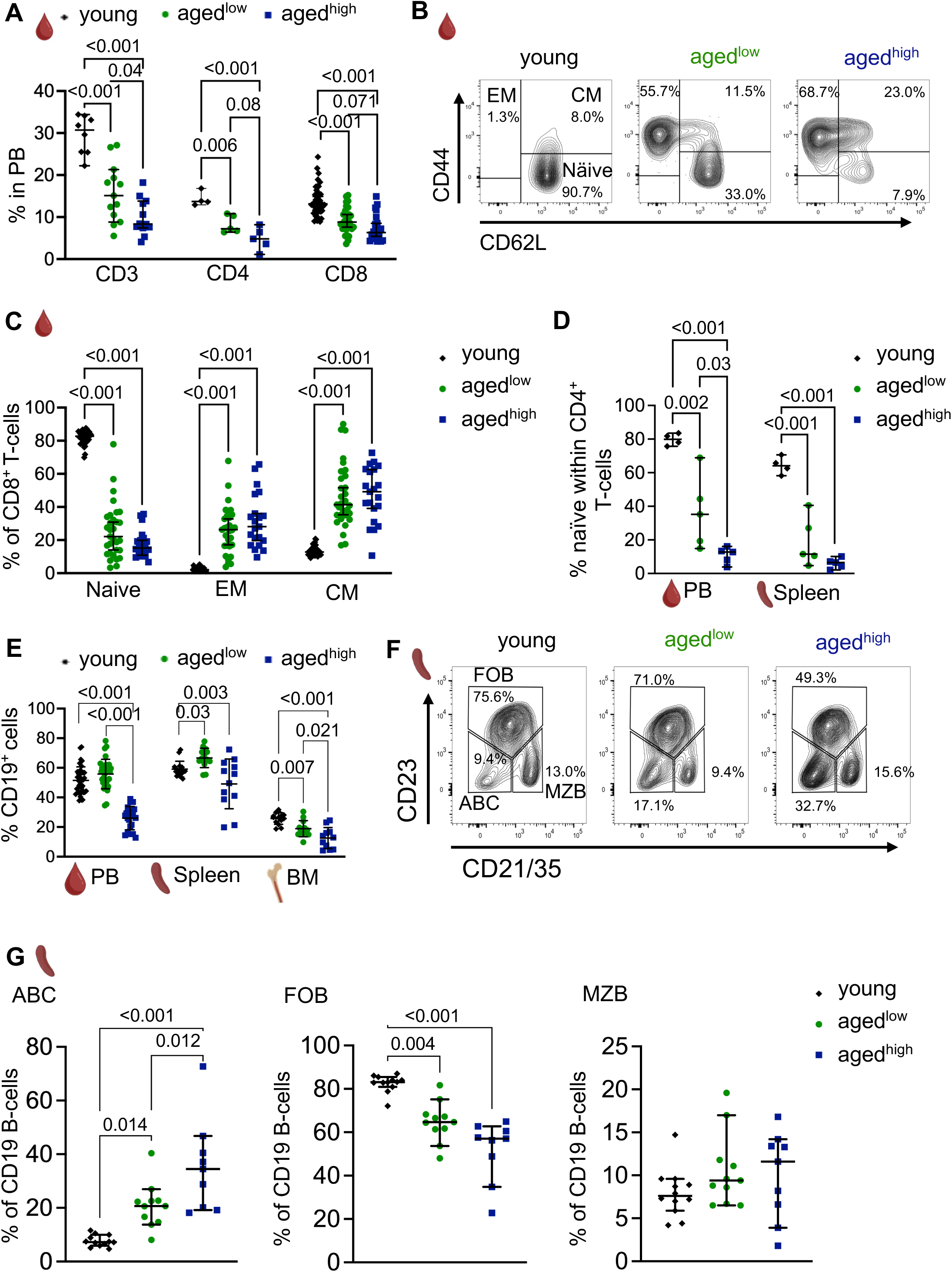
Aged^high^ mice show a more aged adaptive immune system. **A)** Percent of CD3^+^, CD4^+^ and CD8^+^ T-cells in peripheral blood (PB). (CD3: young n = 8; aged^high^ n = 12; aged^low^ n = 13; CD4: young n = 4; aged^low/high^ n = 5; CD8: young n = 40; aged^high^ n = 21; aged^low^ n = 28) **B)** Representative image of the gating strategy for the classification of CD8^+^ T-cell subsets in PB. EM = effector memory; CM = central memory; Naïve = Naïve CD8^+^ T-cells. **C)** Percent of T-cell subsets within CD8^+^ T-cells in PB. EM = effector memory; CM = central memory; Naïve = Naïve CD8 T-cells. (young n = 36-40; aged^high^ n = 21; aged^low^ n = 31) **D)** Percent of naïve CD4^+^ T-cells (CD4^+^, CD62L^+^, CD44^-^) within CD4^+^ T-cells in PB and spleen. (young n = 4; aged^low/high^ n = 5) **E)** Percent of CD19 B-cells in PB, spleen and BM. (PB: young n = 39; aged^high^ n = 21; aged^low^ n = 31; spleen: young n = 16; aged^high^ n = 12; aged^low^ n = 14, BM: young n = 14; aged^high^ n = 11; aged^low^ n = 13) **F)** Gating strategy and **G)** Quantification of CD19 B-cell sub-populations in the spleen. ABC = age-associated B-cells, FOB = follicular B cells, MZB = Marginal-zone B cells. (young n = 12; aged^high^ n = 9; aged^low^ n = 11) Data are presented as medians with 95% CI. Outliers were excluded using the ROUT method (Q = 1%). Statistical analysis was performed using one-way ANOVA with Tukey’s post-hoc test for normally distributed data or Kruskal–Wallis test with Dunn’s multiple-comparison correction for nonparametric data.

Furthermore, also the frequency of B-cells in PB, BM and spleen was significantly reduced in aged^high^ animals compared to aged^low^ and to young animals (**Fig.2E**). We further tested for the accumulation of age-associated B-cells (ABCs) in the spleen^28^ (**Fig.2F**) and found that aged^high^ mice presented with an increase in the frequency of ABCs compared to aged^low^ animals (**Fig.2G**). Collectively, the overall reduction in total T- and B-cell frequencies, the enhanced loss of naïve CD8⁺ and CD4^+^ T-cells and the elevated ABCs in spleen indicate an advanced age of the adaptive immune system in aged^high^ animals.

Fourth, we determined the frequency of distinct types of primitive hematopoietic cells within the BM of young, aged^high^ and aged^low^ animals (**Fig.S4A**). While we observed the well-known age-associated increase in the frequency of long-term HSCs (LT-HSCs) in both age groups, intriguingly, there was no difference in the frequency of LT-HSCs (**Fig.3A**), nor any other type of hematopoietic progenitor cell (HPC), between aged^high^ and aged^low^ animals (**Fig.S4B/C**).

**Figure 3.**
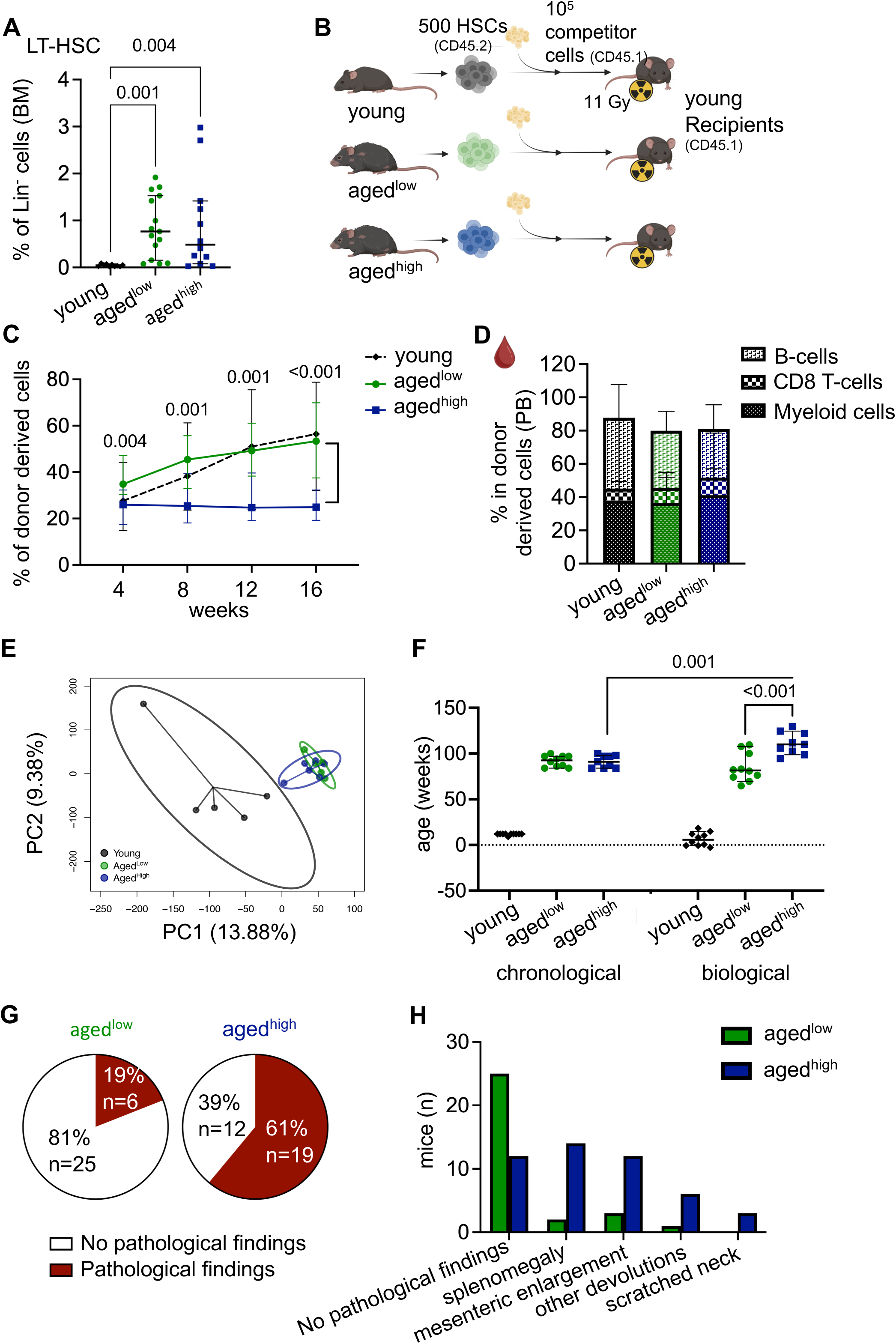
The extent of myeloid skewing correlates with hematopoietic aging, higher biological age and elevated disease burden **A)** Percent of LT-HSCs among lineage negative cells. LT-HSCs = Lin^-^, Sca-1^+^, c-Kit^+^, CD34^low^, CD150^+^, CD48^-^ (young n = 10; aged^high^ n = 12; aged^low^ n = 15). **B)** Transplantation set-up. 500 HSCs (Lin^-^ Sca-1^+^ c-Kit^+^ CD34^low^ Flt3) were isolated from young, aged or aged mice (CD45.2) and transplanted into lethally irradiated recipient mice (CD45.1) together with 10^5^ total bone marrow cells (CD45.1). **C)** Percent of donor derived cells in PB over time (shown in weeks). p-values between aged^high^ and aged^low^ are given, young is significantly different from aged at week 8 and onwards. (young n = 8, age^low/high^d n = 14) **D)** Percent of different cell linages in donor derived cells in PB. (young n = 8, aged n = 14). **E)** Principal component analysis between young, aged and aged HSCs. (young n = 5, aged n = 6, aged n = 5) **F)** Chronological versus biological (predicted) age of young, aged^low^ and aged^high^ mice. Biological age prediction was based on DNA methylation profile of blood cells according to Perez-Correa *et al.* (2022). Aged^high^ mice manifest with an increased biological age compared to aged^low^ mice. Aged^high^ mice 111.1 ± 12.1 weeks to aged^low^ 84.8 ± 15.7 weeks. Significant differences between all aged groups are shown, the young group also differs significantly from both aged groups in terms of chronological and biological age. (young n = 10; aged^high^ n = 9; aged^low^ n = 10). **G)** Terminal macroscopic examination of aged animals; illustrated is the frequency of gross tissue abnormalities in aged^high^ mice (pathological findings 61% in the agedhigh group vs. 19% in the agedlow group, n = 31). **H)** Summary of gross pathological findings. Among 31 animals examined, abnormalities were observed in 6 aged^low^ and 19 aged^high^ mice. Findings were classified into the following categories: splenomegaly (spleen weight > 0.2 g and/or presence of white nodules), mesenteric enlargement (scored as yes/no), other organ devolutions (including pathological changes in the liver, bladder, neck, kidney, or lung such as tumors or cysts), and scratched neck). Individual animals could exhibit more than one abnormality. Data are presented as medians with 95% CI. Outliers were excluded using the ROUT method (Q = 1%). Statistical analysis was performed using one-way ANOVA with Tukey’s post-hoc test for normally distributed data or Welch’s test, for nonparametric data Kruskal–Wallis test with Dunn’s multiple-comparison correction or Mann-Whitney test was applied.

Overall, known aging-associated changes in the frequency of differentiated hematopoietic cells are indeed more pronounced in aged^high^ mice. The extent of myeloid skewing in PB might thus serve as a surrogate marker for the biological age of haematopoiesis in mice.

Next, we assessed HSC function in transplantation experiments (**Fig.3B**). HSCs from aged^high^ animals exhibited a significantly reduced contribution to PB chimerism compared to both HSCs from young and aged^low^ animals (**Fig.3C**). This lower pattern of chimerism was also detected in spleen, BM, and among more differentiated HPCs in BM (**Fig.S5A**). There was no significant difference though among the multiple progenitor population 1 (MPP1) or among LT-HSCs across the groups (**Fig.S5A**).

Surprisingly, HSCs from aged^high^ animals showed within the young recipients no increase in myeloid output in comparison to HSCs from young and aged^low^ animals in PB, BM and spleen (**Fig.3D/S5B**). Comparing the myeloid cell frequencies after transplantation revealed high heterogeneity within all groups with an overall increase in both the young and aged^low^ groups, whereas a decrease was observed in recipients of aged^high^ HSCs (**Fig.S5C**). The transcriptome of HSCs from aged^high^ mice did not imply an intrinsic myeloid bias of aged^high^ HSCs in comparison to aged^low^ HSCs (**Fig.3E/S5D**). Although numerous differentially expressed genes (DEGs) distinguished aged from young HSCs (**Tab.S1**), no DEGs were detected between aged^low^ and aged^high^ HSCs (**Fig.S5D**). Of note, in aged mice, transplantation also restored PB CD8⁺ and naïve T-cell levels and resolved splenic ABC accumulation (**Fig.S5C)**. The reduced function of aged^high^ HSCs upon transplantation might therefore be linked to epigenetic alterations in HSCs rather than being already imprinted in the transcriptome, or it might be a consequence of distinct metabolomic priming of aged^low^ and aged^high^ HSCs. Together, these findings show that HSCs derived from aged^high^ animals are intrinsically less efficient in sustaining hematopoietic reconstitution, while transplantation into young recipients largely resets the high levels of myeloid skewing/differentiation of aged^high^ HSCs.

### Elevated myeloid skewing correlates with higher biological age and higher disease burden

DNA methylation patterns change in an organized and reproducible manner during aging. To investigate whether epigenetic age differs between mice with low or high myeloid bias we applied a targeted epigenetic clock model based on four CG dinucleotides of PB^23^. Despite identical average chronological age (91.7 weeks for aged^low^ and 91.3 weeks for aged^high^), the aged^low^ group exhibited a predicted age of 84.8±15.7 weeks, whereas the aged^high^ group showed a predicted age of 111.1±12.1 weeks, which is a delta of about 26 weeks or 23% (**Fig.3F**). A potential cell type composition bias due to an increased presence of myeloid cells in the aged^high^ mice epigenetic profile, was carefully accounted for during the analysis and excluded as the driver of the difference in predicted age.

Additionally, macroscopic examination at sacrifice showed a markedly higher frequency of disease-related tissue abnormalities in aged^high^ mice (**Fig.3G**). The predominantly observed abnormalities were splenomegaly and mesenteric lymph node enlargement (**Fig.3H/S6**).

In summary, these data strongly suggest that the extent of myeloid skewing in PB is also an indicator for the extent of biological age of an animal, including disease burden.

### The extent of myeloid skewing correlates in older adults with hazard ratio of mortality, morbidity, and mobility

Myeloid skewing has been so far primarily perceived as a hallmark of aging in mice. Thus, we investigated whether the level of myeloid skewing in older adults might also be connected to aspects of biological aging. Thus, we analyzed data from 1,425 individuals aged over 64 years old from the well characterized population-based ActiFE cohort^25,26^. Baseline characteristics of the study population are listed in **Table 1**. Myeloid skewing was quantified by the proportion of myeloid cells—defined as the sum of granulocytes (neutrophils, basophils, eosinophils) and monocytes—relative to the total white blood cell count, mirroring the approach of the determination of the extent of myeloid skewing in our mouse study. Participants were stratified into quartiles based on their extent of myeloid skewing in PB.

Quartile 1 (Q1, low myeloid skewing = older adults^low^) included participants with ≤65.7% myeloid cells, whereas quartile 4 (Q4, high myeloid skewing = older adults^high^) comprised those with ≥77.0% myeloid cells (**Fig.4A**). As expected for a parameter associated with aging there was a linear increase in chronological age of participants from Q1 (73.62±6.59 years) to Q4 (77.82±6.59 years). However, the overall correlation of chronological age and myeloid skewing was modest (r=0.20, **Fig.S7A**). Myeloid skewing in humans was driven by both, a lower frequency of lymphoid cells and an increase of myeloid cells (**Tab.1**).

**Figure 4.**
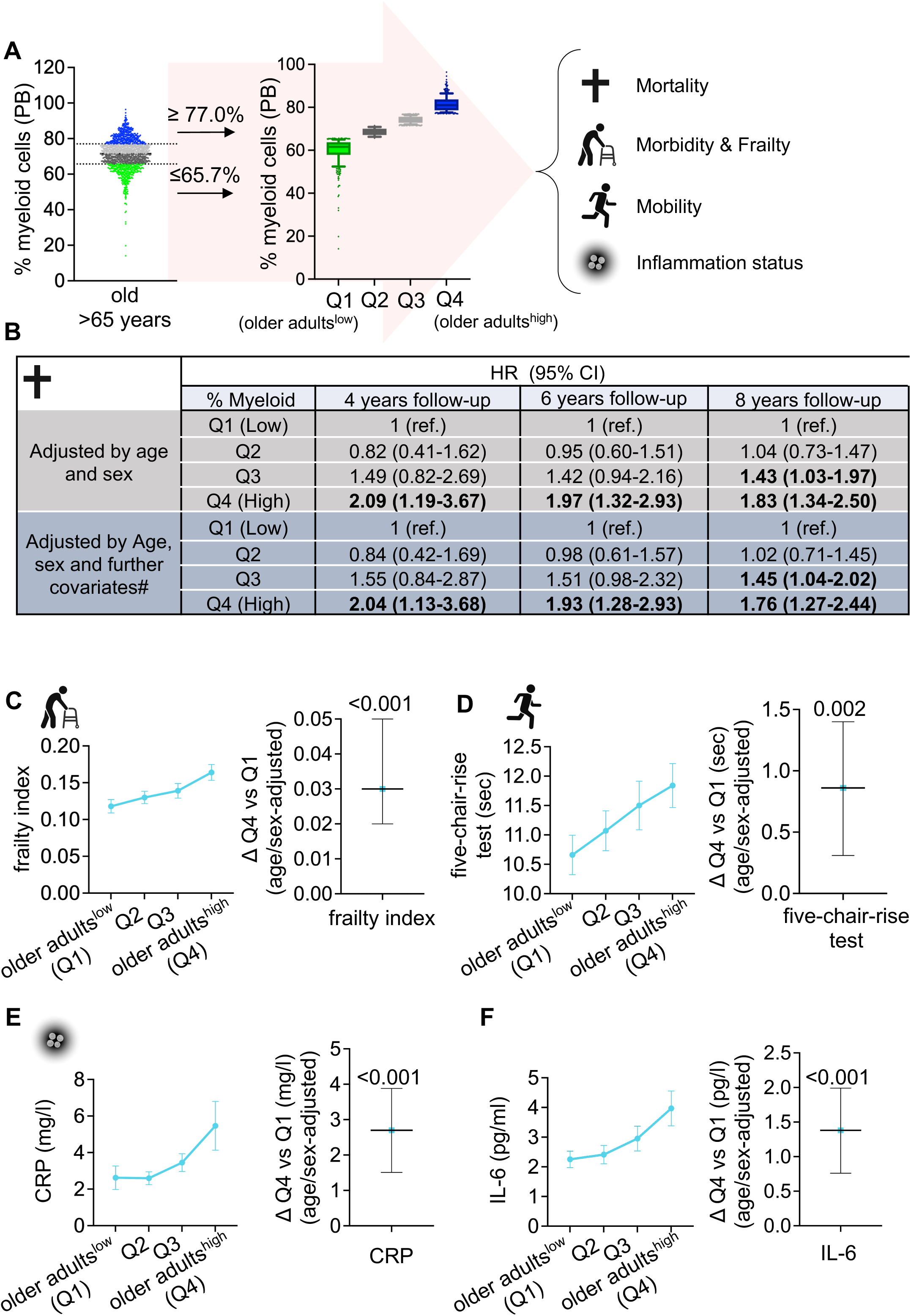
The extent of myeloid skewing correlates in older adults with hazard ratio of mortality, morbidity, and mobility. **A)** Participants (>65 years old) were stratified into quartiles based on the extent of myeloid skewing. Q1 (older adults^low^) represents the 25% of participants with the lowest degree of myeloid skewing, Q4 (older adults^high^) includes the 25% with the highest degree of myeloid skewing. **B)** Association of the percentage of myeloid skewing with hazard ratio for mortality for different time periods of follow-up. Hazard ratios (HR) describe the relative risk of subsequent mortality compared to Q1, older adults^low^ (ref.). Model is adjusted for age and sex, additionally also for other covaries including school education (<=9 / >9 years), BMI (categorised in four categories: underweight, normal, overweight, obese), current smoker, alcohol consumption (daily/less than daily), hypertension, myocardial infarction, cancer, diabetes, number of medications (<5 / >=5); significant changes are bold. **C-F) Left:** Descriptive illustration of specific functional measures and biomarkers within the different quartiles from Q1 older adults^low^ to Q4 older adults^high^ (exact values see Tab.1, unadjusted mean ± SD). **Right:** Adjusted linear regression estimate for Q4 vs Q1 (ref.), controlling for Q2, Q3, age and sex (= differences Q1 older adults^low^ and Q4 older adults^high^). (mean ± SD; p-values derived from t-statistics of the regression coefficients) **C)** Frailty Index based on the 32-item Rockwood scale. (Q1 n=284, Q4 n=285) **D)** Time (in seconds) required to complete five chair rises. (Q1 n=334, Q4 n=328) **E)** C-reactive protein (CRP) concentration (mg/L). (Q1 n=348, Q4 n=353) **F)** Interleukin-6 (IL-6) concentration in serum (pg/mL). (Q1 n=348, Q4 n=354).

First, we determined the extent of the association between percentage of myeloid cells in blood and mortality in follow-up analyses by Cox proportional hazards models (**Fig.4B, Tab.S2**). Older adults^high^ showed consistently a significantly increased risk of mortality compared to older adults^low^ (reference group) (**Fig.4B/Tab.S2**). This association was robust for the different follow-up time points, and was also evident after adjustment for age and sex (**Fig.4B**) and additional covariates (education, BMI, comorbidities, etc.) (**Fig.4B**), although the hazard ratio slightly decreased with duration of follow-up. Second, the number of self-reported doctor diagnosed chronic conditions (hypertension, heart failure, cancer, and myocardial infarction) was highest in older adults^high^ (**Tab.1**). Similarly, the frailty index increased linearly across the quartiles, with the highest level of frailty in older adults^high^ (**Fig.4C/Tab.1**). Even though adjusted differences in frailty index were modest, a 0.03 increase, as seen between older adults^high^ and older adults^low^, equals about one extra health deficit. Third, we assessed the correlation of myeloid skewing to mobility in older adults. Time required to complete the five-chair-rise (assesses functional lower extremity strength, balance, and transitional movement) (**Fig.4D/Tab.1**) as well as measured gait speed (measure for mobility) (**Fig.S7B/Tab.1**) were lowest in older adults^low^ with significant level of increase in older adults^high^. Reference datasets imply that a change of 1 second in the five-chair rise test or a difference of 0,2 seconds in the gait speed are seen in groups of people with 10 years difference in age^29,30^. Similarly, the level of physical activity was highest among older adults^low^ and lowest among older adults^high^ (**Fig.S7C/Tab.1**).

Fourth, we assessed the correlation of the inflammatory markers IL-6 and C-reactive protein (CRP) to the level of myeloid skewing. Older adults^high^ exhibited elevated levels of IL-6 and CRP (**Fig.4E/F/Tab.1**). These results mirror our findings from the association of elevated levels of inflammatory markers in aged^high^ mice and therefore suggest a more general connection between the level of myeloid skewing and the inflammation status. Taken together, older adults^high^ show elevated levels of frailty, higher inflammatory markers, and a strong trend toward reduced physical mobility and muscle function in comparison to older adults^low^. In conclusion, the frequency of myeloid cells in PB in humans correlates with a large set of biomarkers of aging in humans, and is associated with elevated levels of mortality, suggesting that also in humans the frequency of myeloid cells in blood is a marker for biological aging.

In humans, the neutrophil-to-lymphocyte ratio (NLR) is an already established marker of systemic inflammation and a prognostic indicator for various types of diseases^31–33^. In the NLR, the balance of lymphoid cells over neutrophils (NLR) are scored, and for myeloid skewing, the balance of myeloid cells that comprise also neutrophils over lymphoid cells are scored. The NLR therefore resembles in part aspects of the parameter myeloid skewing. Comparison of the associations of myeloid skewing versus the NLR to the difference in limitations and competences in older adults^low^ to adults^high^ indeed revealed that both parameters (myeloid skewing and the NLR) provide very similar differential values with respect to frailty, mobility, inflammation and mortality (**Fig.S7D/Tab.S3**). The marker extent of myeloid skewing in humans is therefore very similar to an already established marker for inflammation/immune senescence/frailty in humans, further validating the frequency of myeloid cells as an additional marker of biological aging in humans. These findings support that the extent of myeloid skewing in mice might indeed serve as a translational biomarker of aging and/or diseases, as this parameter is also associated with biological aging in humans.

## DISCUSSION

Our data reveal that the percentage of myeloid cells in blood in aged mice as well as in older adults might serve as a quantitative biomarker for aging, with higher levels of myeloid skewing associated with multiple markers of advanced aging, disease and an increased risk for mortality in humans. Our data further support that the percentage of myeloid cells in PB in mice is also a robust marker that does not vary significantly depending on how and when the blood sample was harvested. Thus, the extent of myeloid skewing may serve as a convenient and rapidly applicable marker for assessing the hematological/biological aging status in individual experimental mice.

Our murine data also imply that there are likely two distinct types of myeloid skewing during aging: one characterized by a moderate increase in the frequency of myeloid cells without elevated inflammation, and another marked by more pronounced myeloid skewing accompanied by increased levels of inflammatory markers in the blood. While the first seems to be a general consequence of aging, the second seems more closely associated with advanced aging and inflammation. There is a prevailing consensus in the literature that suggests that inflammation is a central driver of the aging process^12,15^. However, our findings indicate that aged mice with lower levels of myeloid skewing—though still higher than in young mice and displaying a “less aged” phenotype compared to aged^high^ mice—still exhibit age-associated functional decline in hematopoiesis. This occurs even in the absence or near absence of elevated inflammatory cytokines. These observations might imply that hallmarks of aging can arise independently of (low) systemic inflammation.

The molecular and cellular mechanisms underlying different levels of myeloid skewing remain speculative, as even the origins of myeloid skewing are still debated. Historically, the emergence of myeloid-biased hematopoiesis from aged HSCs^9,16,34^ has been interpreted as evidence for the clonal expansion of myeloid-biased HSCs with age. Interestingly, recent findings showed that indeed depletion of myeloid-biased HSCs restores a more youth-like immune phenotype^35^. Other studies introduced additional, alternative factors, such as impaired lymphoid hematopoiesis in aged mice^36,37^ or altered interactions between HSCs and BM niche cells^38^. It has also been proposed that the inhibition of lymphocyte maturation outside the BM—in organs such as the thymus and spleen—rather than changes in the intramedullary differentiation environment, may drive myeloid skewing^37^. Here, the fact that HSCs from aged^high^ animals give rise to a rather balanced lineage distribution in young recipient mice suggests that the pronounced myeloid skewing observed upon aging might also be driven by HSCs extrinsic factors, which could be indeed a low-grad systemic inflammation in aged mice. Still, the mechanistic relationship between elevated myeloid cell frequencies, systemic inflammation, and accelerated aging remains speculative and warrants further investigation. Additional experiments are also required to determine whether elevated levels of myeloid skewing are a driver or an indicator of an already underlying inflammation and disease.

The extent of myeloid skewing in humans parallels the NLR, both conceptually and in their association with biological markers of aging. NLR is a well-established clinical indicator of systemic inflammation, immune status, and disease severity, and is has been also used in aging research^31–33^. Similarly, myeloid skewing reflects age-related immune alterations and health decline, making it a promising and clinically relevant marker in murine models. This highlights myeloid skewing as a useful marker of biological aging in chronologically aged mice, and the alignment with human cohort data supports translational validity of murine data. Incorporating quantitative determination of the level of myeloid skewing as a standard assessment could therefore enhance the rigor and interpretability of studies on aging and age-related disease in mice, but also in humans.

In conclusion, our data suggest that the extent of myeloid skewing in blood is a biological marker of aging for both mice and humans. Given that myeloid skewing is easily quantifiable and exhibits in mice consistent results regardless of sampling technique or time, it is well-suited to establish an understanding of biological heterogeneity in chronologically aged mice. Intriguingly, it may also serve as an additional longitudinal marker that may assist monitoring of functional decline in older adults.

## Supporting information

Full Supplementary

