## Supplementary material for "The extent of myeloid skewing in blood is a biomarker of biological aging in mice and humans": Full Supplementary

**Supplementary Information**

**MATERIALS**

***Table S1:*** *List of antibodies used in this study.*

| Antibody | Fluorochrome | Company | Cat.-Nr- | Clone | LOT |
|---|---|---|---|---|---|
| CD16/32 (Fc-Block) | - | eBioscience | 14-0161-85 | 93 | 2334894 |
| CD45.2 | APC-Vio770 | Miltenyi Biotec | 130-119-128 | 104-2 | 5231003922 |
| | PerCp-Cy5.5 | eBioscience | 45-0454-82 | 104 | 2331111 |
| Ly6G/C (Gr-1) | eFluor450 | eBioscience | 48-5931-82 | RB6-8C5 | 2422188 |
| CD11b | AF700 | eBioscience | 56-0112-82 | M1/70 | 2433037 |
| CD3e | PE-Cy7 | Thermo Fisher | 25-0031-82 | 145-2C11 | 2619646 |
| CD8a | PE-Cy7 | Thermo Fisher | 25-0081-82 | 53-6.7 | 2331993 |
| | APC | Thermo Fisher | 17-0081-82 | 53-6.7 | 4321418 |
| CD4 | PerCp-Cy5.5 | Thermo Fisher | 45-0042-82 | RM4-5 | 2286404 |
| CD19 | APC | eBioscience | 17-0193-82 | 1D3 | 2514154 |
| CD49d | PE | BioLegend | 103607 | R1-2 | B363406 |
| CD44 | FITC | BD Pharmingen | 553133 | IM7 | 1109185 |
| CD62L | PerCp-Cy5.5 | Thermo Fisher | 45-0621-82 | MEL-14 | 2480285 |
| CD23 | PE-Cy7 | eBioscience | 25-0232-82 | B3B4 | 1981960 |
| CD21/35 | eFluor450 | BioLegend | 123414 | 7E9 | B316532 |
| Lineage | Biotin | Miltenyi Biotec | 130-090-858 | - | 5230705797 |
| Streptavidin | eFluor450 | eBioscience | 48-4317-82 | - | 2527387 |
| c-Kit | AF700 | eBioscience | 56-1172-82 | ACK2 | 2505081 |
| Ly6A/E (Sca-1) | PE-Cy7 | eBioscience | 25-5981-82 | D7 | 2514094 |
| CD34 | APC | eBioscience | 50-0341-82 | RAM34 | 2305277 |
| | FITC | eBioscience | 11-0341-82 | RAM34 | 2310844 |
| CD150 | PE | BioLegend | 115904 | TC15-12F12.2 | B367830 |
| CD48 | AF647/APC | BioLegend | 103416 | HM48-1 | B379738 |
| CD135 (Flt-3) | PE | Thermo Fisher | 12-1351-83 | A2F10 | 2088477 |
| IL7Ra | APC-Cy7 | eBioscience | 47-1271-82 | A7R34 | 2611853 |
| CD16/32 | PE | eBioscience | 12-0161-82 | 93 | 2422285 |

**SUPPLEMENTARY FIGURE LEGENDS**

**Figure S1.**

**A)** Correlation between myeloid cell percentage in PB measured by flow cytometry and the % of granulocytes obtained from the blood cell counter ScilVet. (n = 207, Pearson correlation, different bleeding methods). **B)** Blood cell counts between young (10-16 weeks) and aged (> 80 weeks) mice measured with ScilVet. Myeloid cells contain number of monocytes, granulocytes and eosinophiles. (young n = 28-29, aged n = 50-53). **C)** Analysis of myeloid percentage in PB with different bleeding methods. (n = 8-25). **D)** % of myeloid cells in PB of young and aged mice bled at 10 am and 10 pm. (young n = 18, aged n = 28). Data are presented as medians with 95% CI. Outliers were excluded using the ROUT method (Q = 1%). Statistical analysis was performed using non-parametric Mann-Whitney test or Welch´s test between young and old.

**Figure S2.**

**A)** % of myeloid cells ($Mac1^{+}$ and/or $Gr1^{+}$) in bone marrow (BM) and spleen. (BM: young n = 14; $aged^{high}$ n = 11; $aged^{low}$ n = 13; Spleen: young n = 16; $aged^{high}$ n = 12; $aged^{low}$ n = 15). Data are presented as medians with 95% CI, no outliers were excluded and statistical analysis was performed using one-way ANOVA with Tukey's post-hoc test. **B/C)** Spearman correlation was performed between % of myeloid cells in PB and serum concentrations of the different cytokines for aged mice. r-values are displayed, and there was no significant difference observed **C)** Rantes and IL-6 and **D)** IL-10 and MCP-1 (n=14).

**Figure S3.**

**A/B)** % of $CD3^{+}$, $CD4^{+}$, $CD8^{+}$ T-cells **A)** in spleen **B)** in BM. (CD3: young n = 5; $aged^{low/high}$ n = 6; CD4: young n = 4; $aged^{low/high}$ n = 5; CD8: young n = 14-16; $aged^{low}$ n = 13-15; $aged^{high}$ n = 11) **C)** % of T-cell subsets in $CD8^{+}$ T-cells in spleen. EM = effector memory; CM = central memory; Naïve = Naïve $CD8^{+}$ T-cells. (young n = 14; $aged^{low}$ n = 11-13; $aged^{high}$ n = 11) **D)** % of T-cell subsets in $CD8^{+}$ T-cells in BM. EM = effector memory; CM = central memory; Naïve = Naïve CD8 T-cells. (young n = 17-20; $aged^{low}$ n = 20; $aged^{high}$ n = 16-17). Data are presented as medians with 95% CI. Outliers were excluded using the ROUT method (Q = 1%). Statistical analysis was performed using one-way ANOVA with Tukey's post-hoc test for normally distributed data or Kruskal–Wallis test with Dunn's multiple-comparison correction for nonparametric data.

**Figure S4.**

**A)** Gating strategy and marker panel for the different hematopoietic stem and progenitor cell populations. **B)** % of the different multipotent progenitor (MPP) populations (MPP1-4) in linage negative cells. (young n = 10; aged$^{high}$ n = 10-12; aged$^{low}$ n = 15; median with 95% CI, one-way ANOVA). **C)** % of the more differentiated progenitor cell populations in linage negative cells; CLP = common lymphoid progenitor, GMP = granulocyte-monocyte progenitor, CMP = common myeloid progenitor, MEP = megakaryocytic-erythroid progenitor). (young n = 7; aged$^{high}$ n = 8; aged$^{low}$ n = 9; median with 95% CI, one-way ANOVA). Data are presented as medians with 95% CI. Outliers were excluded using the ROUT method (Q = 1%). Statistical analysis was performed using one-way ANOVA with Tukey's post-hoc test for normally distributed data or Kruskal–Wallis test with Dunn's multiple-comparison correction for nonparametric data.

**Figure S5.**

**A)** % of donor derived cells within different hematopoietic tissues and hematopoietic stem and progenitor cells 16 weeks after transplantation. (young n = 8; aged$^{low/high}$ n = 14) **B)** % of different cell linages in donor derived cells in BM and spleen. (young n = 8, aged$^{low/high}$ n = 14) **C)** % of different cell types in PB and spleen in donors (before TXT, red dots) and in recipients (after transplantation, black dots). (young n = 3 donors (2-3 mice pooled per n), 8 recipients; aged$^{low/high}$ n = 2-6 donors, 6-14 recipients) **D)** Heatmap of log2 fold change of DEGs from RNA-Sequencing of young, aged$^{low}$ and aged$^{high}$ mice with a phylogenetic depiction of sample similarity (young n = 5, aged$^{low}$ n = 5, aged$^{high}$ n = 6). Data are presented as medians with 95% CI. Outliers were excluded using the ROUT method (Q = 1%). Statistical analysis was performed using one-way ANOVA with Tukey's post-hoc test for normally distributed data or Kruskal–Wallis test with Dunn's multiple-comparison correction for nonparametric data.

**Figure S6.**

**A)** Representative images of mesenteric lymph nodes categorized as normal or showing mesenteric volume enlargement. **B) Left:** Representative spleens from young, aged$^{low}$, and aged$^{high}$ animals. **Right:** Quantification of spleen weights. (young n = 12; aged$^{low/high}$ n = 16; medians with 95% CI; Kruskal–Wallis test with Dunn's multiple-comparison correction; outliers were not removed)

**Figure S7.**

**A)** Spearman correlation between age (in years) at the time of the visit and the % of myeloid skewing in peripheral blood. (n=1425) **B-D) Left:** Descriptive illustration of specific functional measures and biomarkers within the different quartiles from Q1 older adults$^{low}$ to Q4 older adults$^{high}$. (exact values see Tab.1, unadjusted mean ± SD) **Right:** Adjusted linear regression estimate for Q4 vs Q1 (ref.), controlling for Q2, Q3, age and sex (= differences Q1 older adults$^{low}$ and Q4 older adults$^{high}$). (mean ± SD; p-values derived from t-statistics of the regression coefficients) **B)** Gait speed test, fastest time (in seconds) of the two runs **C)** Daily mean of walking in minutes **D)** Comparison of the percentage of myeloid skewing and the neutrophil-to-lymphocyte ratio (NLR), both stratified into quartiles. Shown is adjusted linear regression estimate for Q4 vs Q1 (ref.), controlling for Q2, Q3, age and sex (= differences Q1 older adults$^{low}$ and Q4 older adults$^{high}$). (mean ± SD; p-values derived from t-statistics of the regression coefficients).

### Supplementary Figures

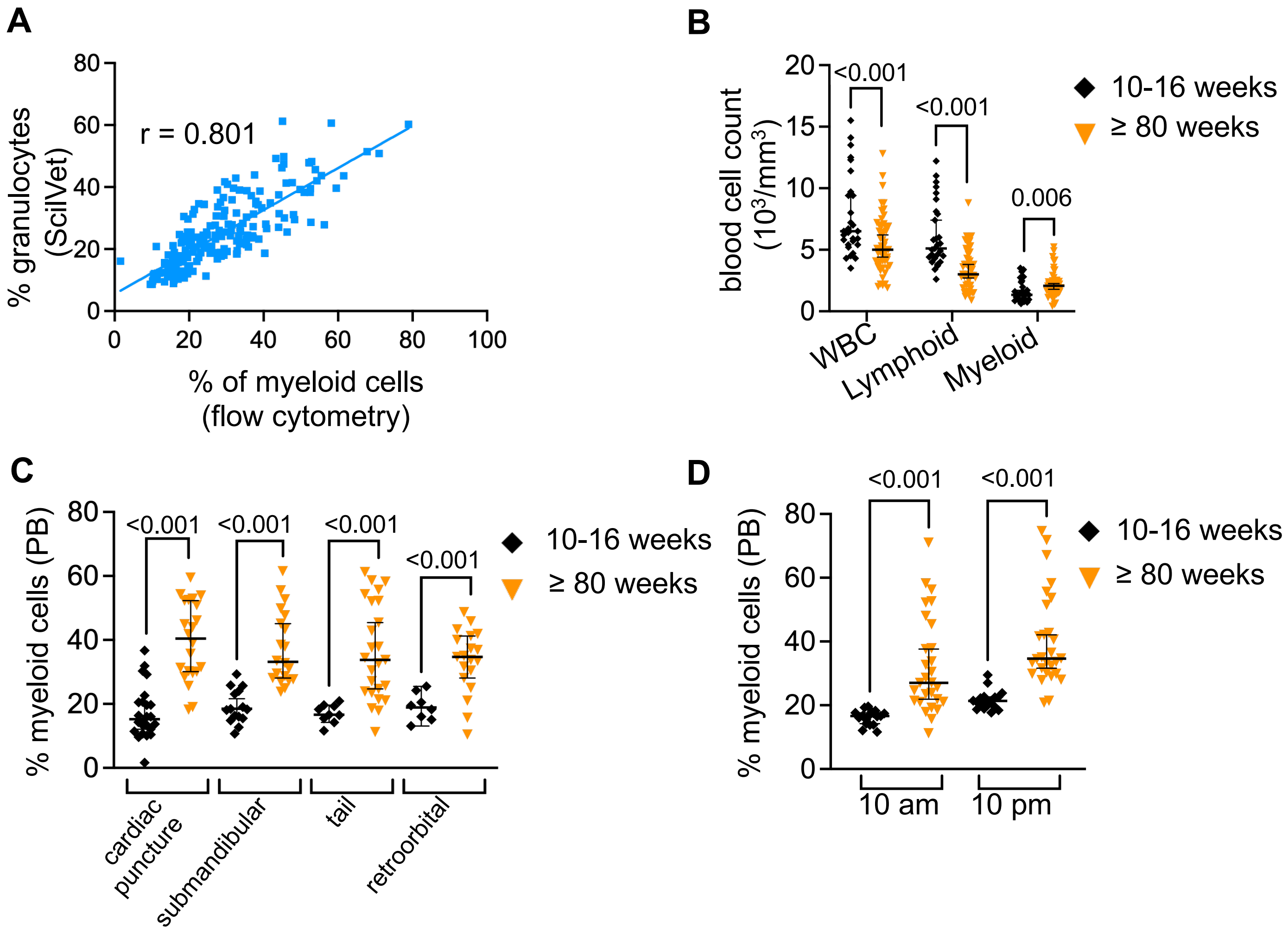
A
r = 0.801
% granulocytes (ScilVet)
% of myeloid cells (flow cytometry)
B
blood cell count ($10^3$/mm$^3$)
<0.001
<0.001
0.006
WBC
Lymphoid
Myeloid
10-16 weeks
≥ 80 weeks
C
% myeloid cells (PB)
<0.001
<0.001
<0.001
<0.001
cardiac puncture
submandibular
tail
retroorbital
10-16 weeks
≥ 80 weeks
D
% myeloid cells (PB)
<0.001
<0.001
10 am
10 pm
10-16 weeks
≥ 80 weeks

**Figure S1**

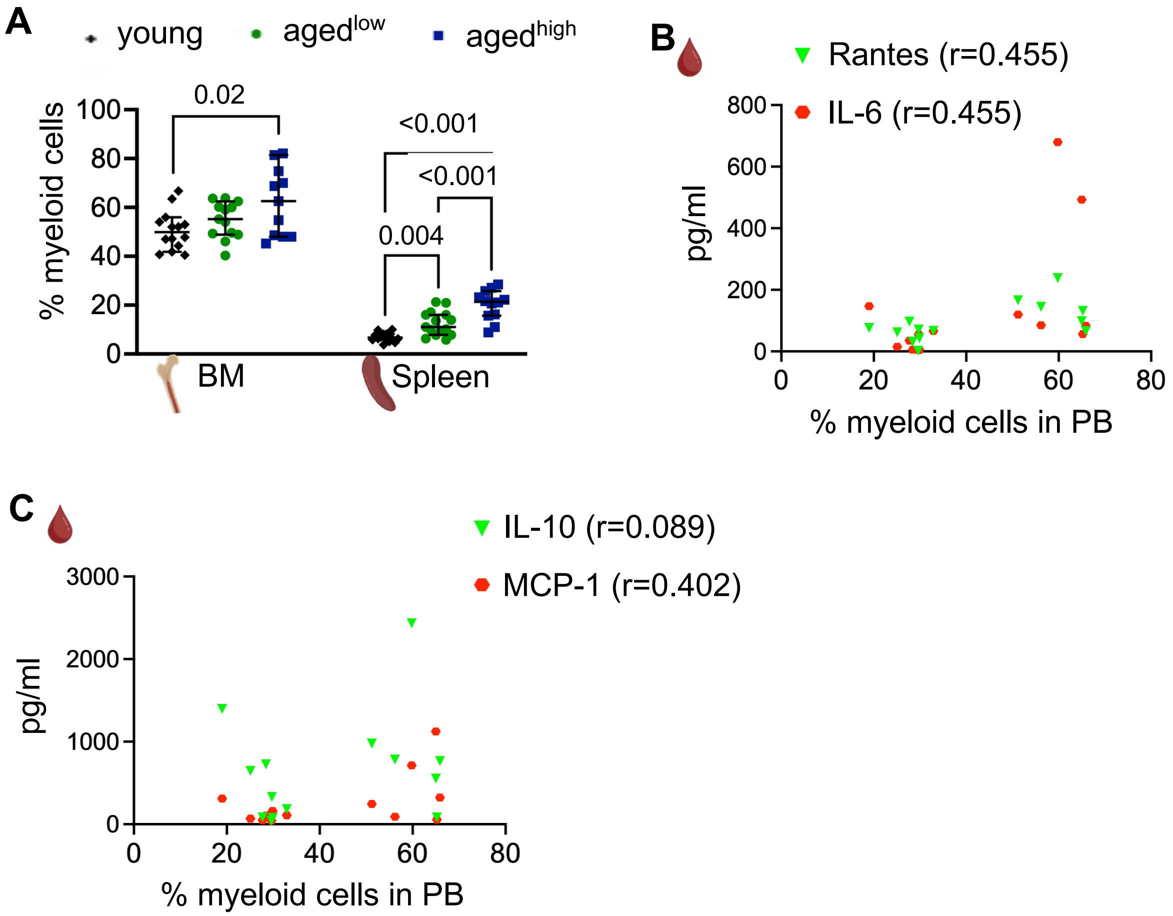

A
young
aged^low
aged^high
% myeloid cells
0.02
<0.001
<0.001
0.004
BM
Spleen
B
Rantes (r=0.455)
IL-6 (r=0.455)
pg/ml
% myeloid cells in PB
C
IL-10 (r=0.089)
MCP-1 (r=0.402)
pg/ml
% myeloid cells in PB


**Figure S2**

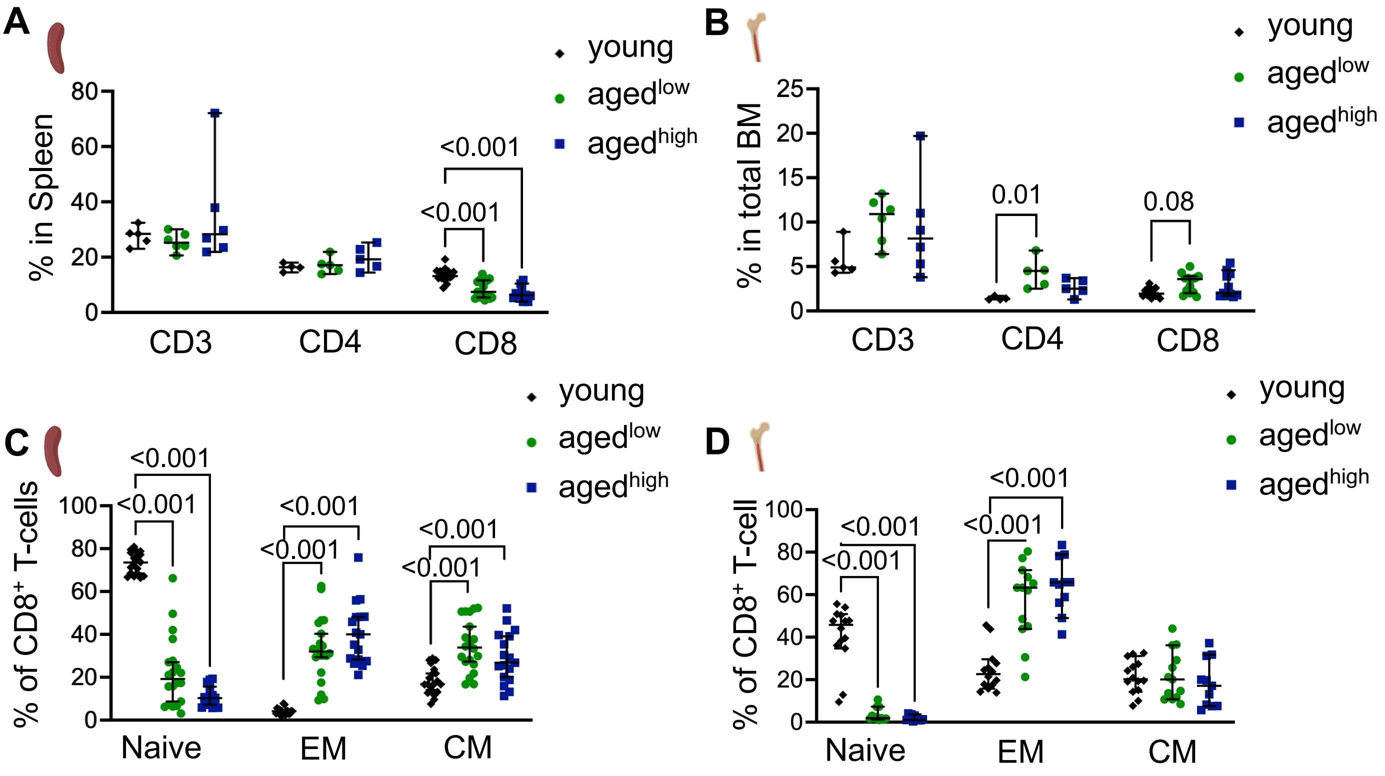
A
young
aged^low
aged^high
% in Spleen
80
60
40
20
0
<0.001
<0.001
CD3
CD4
CD8
B
young
aged^low
aged^high
% in total BM
25
20
15
10
5
0
0.01
0.08
CD3
CD4
CD8
C
young
aged^low
aged^high
% of CD8+ T-cells
100
80
60
40
20
0
<0.001
<0.001
<0.001
<0.001
<0.001
<0.001
Naive
EM
CM
D
young
aged^low
aged^high
% of CD8+ T-cell
100
80
60
40
20
0
<0.001
<0.001
<0.001
<0.001
Naive
EM
CM

**Figure S3**

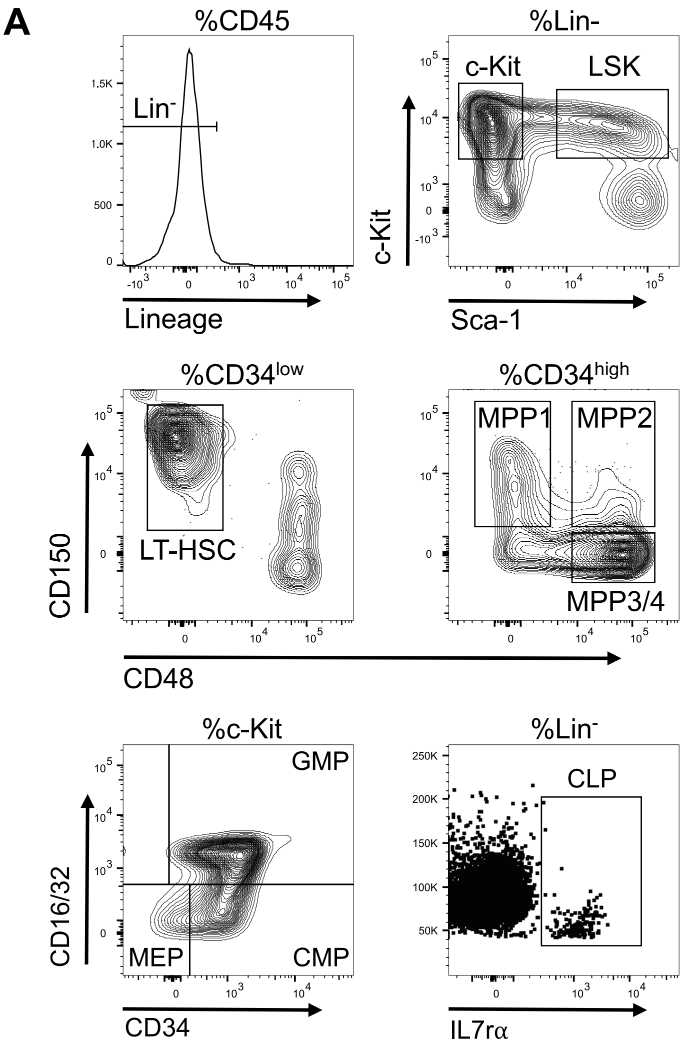


| Cell Type | Surface Marker |
|---|---|
| LT-HSC | LSK CD34$^{low}$ CD150$^{+}$ CD48$^{-}$ |
| MPP1 | LSK CD34$^{high}$ CD150$^{+}$ CD48$^{-}$ |
| MPP2 | LSK CD34$^{high}$ CD150$^{+}$ CD48$^{+}$ |
| MPP3/4 | LSK CD34$^{high}$ CD150$^{-}$ CD48$^{+}$ |
| MEP | Lin$^{-}$ c-Kit$^{+}$ Sca1$^{-}$CD34$^{-}$ CD16/32$^{-}$ |
| CMP | Lin$^{-}$ c-Kit$^{+}$ Sca1$^{-}$CD34$^{+}$ CD16/32$^{-}$ |
| GMP | Lin$^{-}$ c-Kit$^{+}$ Sca1$^{-}$CD34$^{+}$ CD16/32$^{+}$ |
| CLP | Lin$^{-}$ IL7rα$^{+}$ |

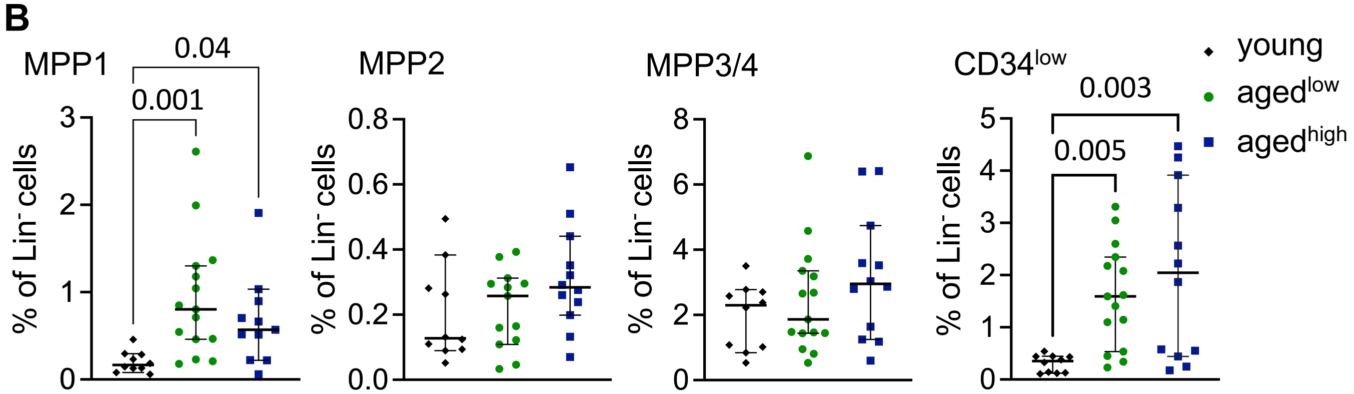


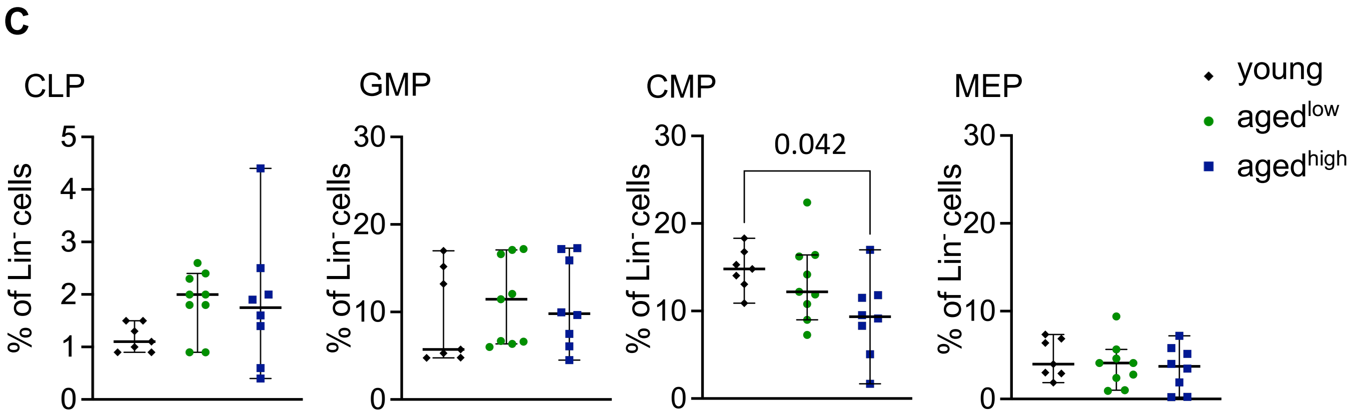


**Figure S4**

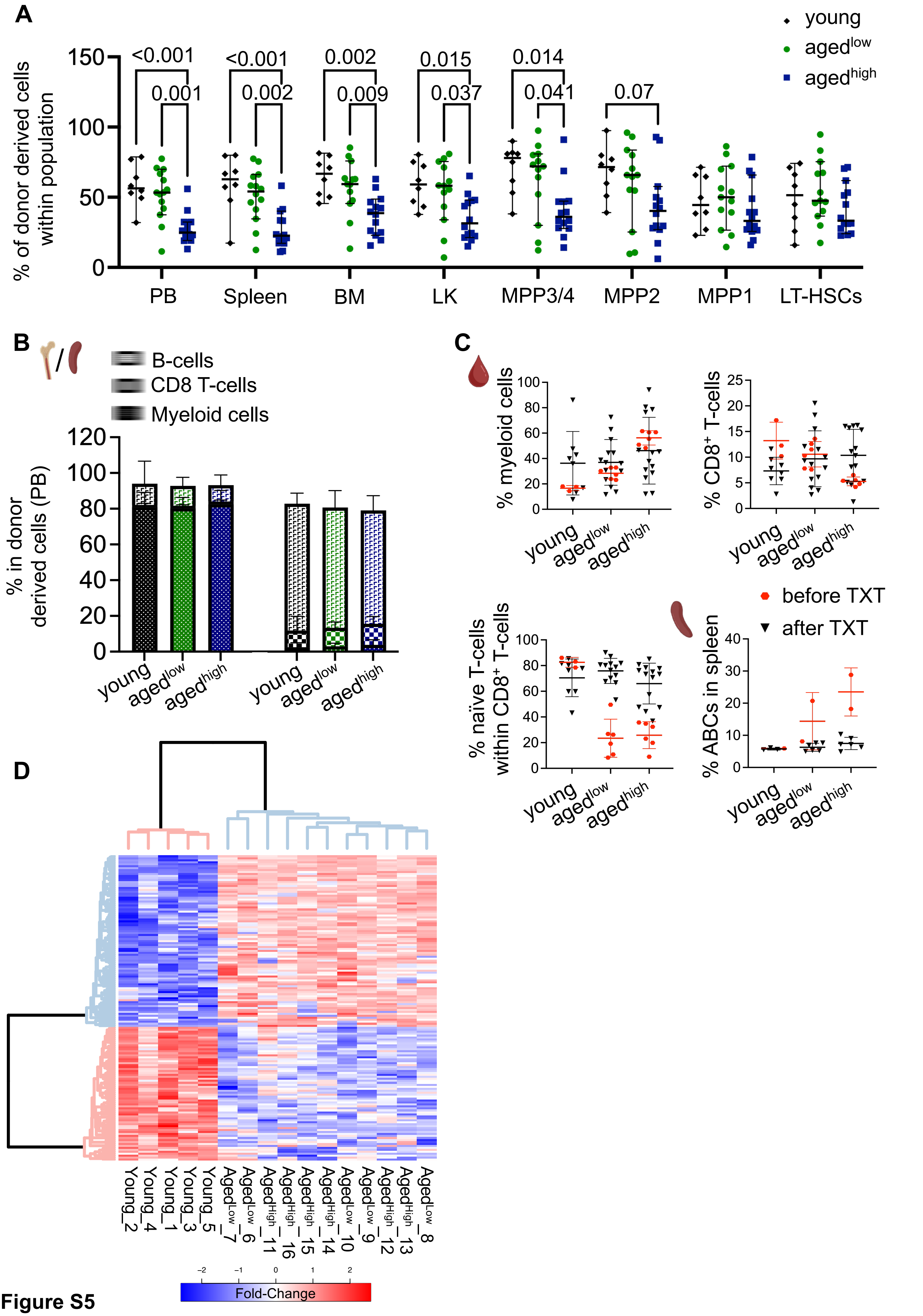
A
young
aged^low
aged^high
% of donor derived cells within population
150
100
50
0
<0.001
0.001
<0.001
0.002
0.002
0.009
0.015
0.037
0.014
0.041
0.07
PB
Spleen
BM
LK
MPP3/4
MPP2
MPP1
LT-HSCs
B
B-cells
CD8 T-cells
Myeloid cells
% in donor derived cells (PB)
120
100
80
60
40
20
0
young
aged^low
aged^high
young
aged^low
aged^high
C
% myeloid cells
% CD8+ T-cells
young
aged^low
aged^high
before TXT
after TXT
% naïve T-cells within CD8+ T-cells
% ABCs in spleen
D
Young_2
Young_4
Young_1
Young_3
Young_5
Aged^Low_7
Aged^Low_6
Aged^High_11
Aged^High_16
Aged^High_15
Aged^High_14
Aged^Low_10
Aged^Low_9
Aged^High_12
Aged^High_13
Aged^Low_8
−2
−1
0
1
2
Fold-Change

Figure S5

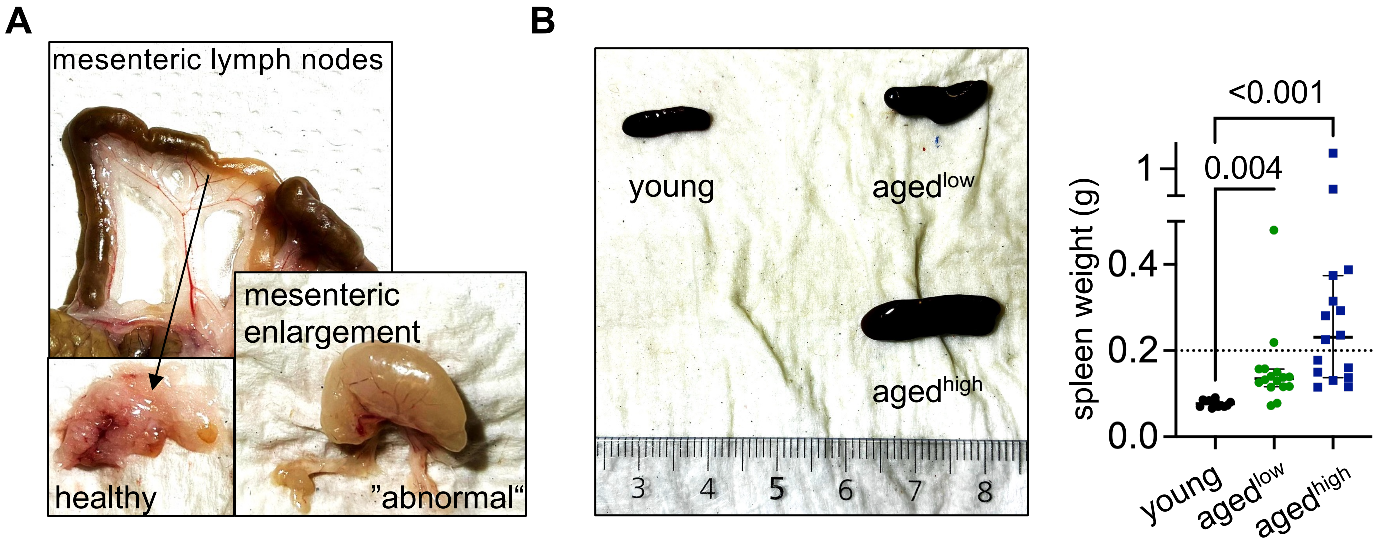
A
mesenteric lymph nodes
mesenteric
enlargement
healthy
"abnormal"
B
young
aged^low
aged^high
3
4
5
6
7
8
<0.001
0.004
1
0.4
0.2
0.0
spleen weight (g)
young
aged^low
aged^high

**Figure S6**

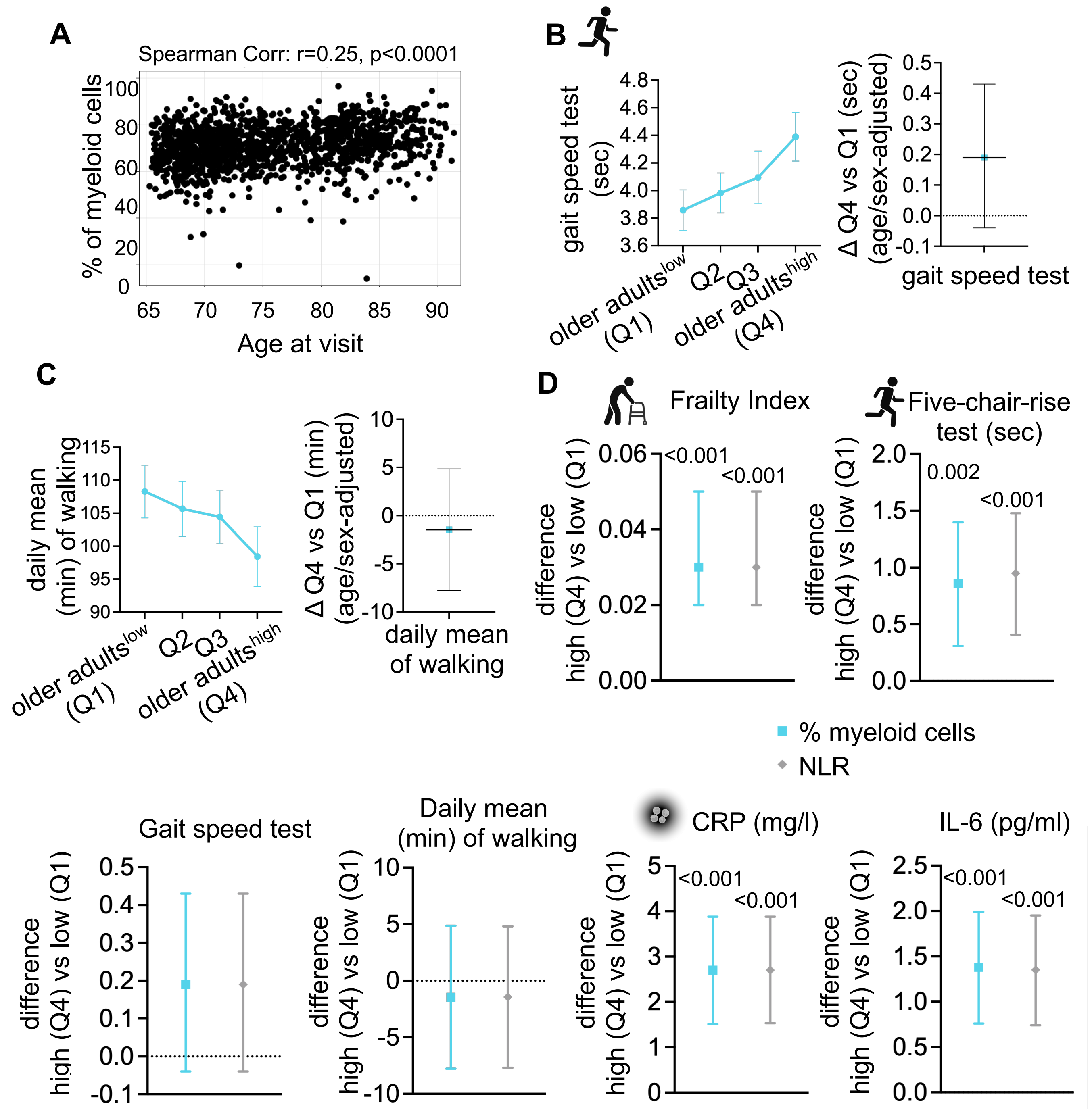
A
Spearman Corr: r=0.25, p<0.0001
% of myeloid cells
Age at visit
B
gait speed test (sec)
older adults^low (Q1)
Q2
Q3
older adults^high (Q4)
Δ Q4 vs Q1 (sec) (age/sex-adjusted)
gait speed test
C
daily mean (min) of walking
older adults^low (Q1)
Q2
Q3
older adults^high (Q4)
Δ Q4 vs Q1 (min) (age/sex-adjusted)
daily mean of walking
D
Frailty Index
difference high (Q4) vs low (Q1)
<0.001
<0.001
Five-chair-rise test (sec)
difference high (Q4) vs low (Q1)
0.002
<0.001
% myeloid cells
NLR
Gait speed test
difference high (Q4) vs low (Q1)
Daily mean (min) of walking
difference high (Q4) vs low (Q1)
CRP (mg/l)
difference high (Q4) vs low (Q1)
<0.001
<0.001
IL-6 (pg/ml)
difference high (Q4) vs low (Q1)
<0.001
<0.001

**Figure S7**

### Supplementary Tables

**Table S1**

Shared differentially expressed genes (DEG) for agedlow and agedhigh in comparison to young HSCs analysed from RNA Sequencing data. Shown are downregulated (blue) and upregulated (red) DEGs shared for both (aged$^{low}$,aged$^{high}$) as well as one of them only.

| DOWN in BOTH | DOWN in aged-low ONLY | DOWN in aged-high ONLY |
|---|---|---|
| Dock10 | Ncoa2 | Rtel1 |
| Arl4c | Ikzf2 | Tspan2 |
| Sell | Inafm2 | Med13l |
| Arhgap30 | Prune1 | Zan |
| Cd34 | Bcl9 | Tmsb10 |
| Arhgap15 | Tmem64 | Rnf225 |
| Itga4 | Macf1 | Dnase1l3 |
| Cntln | Cnot6l | Rfx7 |
| Ptprf | Git2 | Ikzf1 |
| Lat2 | Hip1r | Arl5c |
| Tmem121b | Flt3 | Mtss1 |
| Mgst1 | Fbxl14 | Csf2rb |
| Tm6sf1 | Etnk1 | H2-Ob |
| Tespa1 | Ccer2 | Tnf |
| Vmn2r84 | Gramd1a | |
| Irf2bp2 | Ankrd27 | |
| Ccl3 | Fchsd2 | |
| Rnf144a | Slc35d3 | |
| Runx2 | Amd2 | |
| Satb1 | Gnptab | |
| Lrg1 | Hmga2 | |
| Bin1 | Chd9 | |
| Zfp608 | Fam117a | |
| Pcgf5 | Rreb1 | |
| | Macroh2a1 | |
| | Yy1 | |
| | Nrip1 | |
| | Rock1 | |
| | Mc5r | |
| | Map4k2 | |

| UP in BOTH | UP in aged-low ONLY | UP in aged-high ONLY |
|---|---|---|
| Kcnb2 | Cxcr1 | Creb3l1 |
| Selp | Chpf | Exd1 |
| Slamf1 | Epha4 | Ptpn13 |
| Fap | Npdc1 | Rbm19 |
| Serping1 | Spint1 | Gm49342 |
| Matn4 | Zbp1 | Esyt3 |
| Rbpjl | Chil5 | Sox30 |
| Dok5 | Npnt | Zmynd15 |
| Tm4sf1 | Slc31a1 | Mboat1 |
| Mab21l2 | Gbp9 | Dsp |
| Muc1 | Kpna7 | Nxnl2 |
| Gstm2 | Tmem176a | Trappc12 |
| Clca3a1 | Gm44511 | Arl2 |
| Gipc2 | Plekhb1 | |
| Ptpru | Il18bp | |
| Asns | Fam241b | |
| Lmcd1 | Igtp | |
| Ntf3 | Aldh3a1 | |
| Klrb1c | Aspa | |
| Clec1a | Ly6e | |
| Klra17 | Alg3 | |
| Erp27 | H2-Aa | |
| Nupr1 | H2-Eb1 | |
| Sult1a1 | Ehd3 | |
| Zg16 | Kcnn2 | |
| Eya4 | Batf2 | |
| Mt2 | | |
| Mt1 | | |
| Pnp | | |
| Phf11d | | |
| Clu | | |
| Plscr2 | | |
| Traip | | |
| Uba7 | | |
| Gm12185 | | |
| Ifi47 | | |
| Rcvrn | | |
| Trim47 | | |
| C7 | | |
| Arhgap8 | | |
| Cpne8 | | |
| Smagp | | |
| Jam2 | | |
| Meiob | | |
| Enpp5 | | |
| Rorb | | |
| Aldh1a1 | | |
| Pdlim1 | | |

**Table S2**

Association of the % of myeloid skewing with mortality for different time periods of follow-up. Hazard ratios describe the relative change in mortality compared to Q1 low (ref.). Model is unadjusted, significant changes are bold.

| | HR (95% CI) | | | |
|---|---|---|---|---|
| | % Myeloid cells | 4 years follow-up | 6 years follow-up | 8 years follow-up |
| Crude | Q1 (Low) | 1 (ref.) | 1 (ref.) | 1 (ref.) |
| | Q2 | 1.05 (0.53-2.07) | 1.23 (0.78-1.95) | 1.35 (0.96-1.91) |
| | Q3 | **2.20 (1.22-3.96)** | **2.13 (1.41-3.23)** | **2.10 (1.53-2.90)** |
| | Q4 (High) | **4.06 (2.34-7.04)** | **3.79 (2.57-5.60)** | **3.38 (2.49-4.59)** |

**Table S3**

Association of the neutrophil to lymphoid ratio (NLR) with mortality for different time periods of follow-up. Hazard ratios describe the relative change in mortality compared to Q1 low (ref.). Models are adjusted for age and sex, additionally also for other covaries including school education (<=9 / >9 years), BMI (categorised in four categories: underweight, normal, overweight, obese), current smoker, alcohol consumption (daily / less than daily), hypertension, myocardial infarction, cancer, diabetes, number of medications (<5 / >=5), significant changes are bold.

| | | **HR (95% CI)** | | |
|---|---|---|---|---|
| | | 4 years follow-up<br>N= 129 deceased | 6 years follow-up<br>N= 254 deceased | 8 years follow-up<br>N= 397 deceased |
| | | NLR | NLR | NLR |
| **Crude** | older adults$^{low}$ (Q1) | 1 (ref.) | 1 (ref.) | 1 (ref.) |
| | Q2 | 1.01 (0.52-1.98) | 1.09 (0.70-1.71) | 1.17 (0.84-1.64) |
| | Q3 | **1.90 (1.06-3.43)** | **1.81 (1.21-2.71)** | **1.80 (1.32-2.46)** |
| | older adults$^{high}$ (Q4) | **4.07 (2.38-6.96)** | **3.50 (2.41-5.07)** | **3.08 (2.30-4.12)** |
| Adjusted by age and sex | older adults$^{low}$ (Q1) | 1 (ref.) | 1 (ref.) | 1 (ref.) |
| | Q2 | 0.85 (0.43-1.66) | 0.89 (0.57-1.40) | 0.95 (0.68-1.33) |
| | Q3 | 1.36 (0.75-2.46) | 1.25 (0.83-1.88) | 1.26 (0.92-1.73) |
| | older adults$^{high}$ (Q4) | **2.14 (1.24-3.70)** | **1.83 (1.25-2.68)** | **1.69 (1.25-2.27)** |
| Adjusted by Age, sex and further covariates# | older adults$^{low}$ (Q1) | 1 (ref.) | 1 (ref.) | 1 (ref.) |
| | Q2 | 0.82 (0.41-1.64) | 0.88 (0.56-1.40) | 0.89 (0.63-1.26) |
| | Q3 | 1.38 (0.75-2.55) | 1.28 (0.84-1.94) | 1.28 (0.93-1.76) |
| | older adults$^{high}$ (Q4) | **2.04 (1.15-3.61**) | **1.75 (1.18-2.60)** | **1.60 (1.17-2.18)** |

**Table S3**
